# Comparing mouse and human cingulate cortex organization using functional connectivity

**DOI:** 10.1101/2023.09.04.556193

**Authors:** Aran T.B. van Hout, Sabrina van Heukelum, Matthew F.S. Rushworth, Joanes Grandjean, Rogier B. Mars

## Abstract

The subdivisions of the extended cingulate cortex of the human brain are both implicated in a number of high-level behaviors and affected by a range of neuropsychiatric disorders. Its anatomy, function, and response to therapeutics are often studied using non-human animals, including the mouse. However, the similarity of human and mouse frontal cortex, including cingulate areas, is still not fully understood. Some accounts emphasize resemblances between mouse cingulate cortex and human cingulate cortex while others emphasize similarities with human granular prefrontal cortex. We use comparative neuroimaging to study the connectivity of the cingulate cortex in the mouse and human, allowing comparisons between mouse ‘gold standard’ tracer and imaging data, and, in addition, comparison between the mouse and the human using comparable imaging data. We find overall similarities in organization of the cingulate between species, including anterior and midcingulate areas and a retrosplenial area. However, human cingulate contains subareas with a more fine-grained organization than is apparent in the mouse and it has connections to prefrontal areas not present in the mouse. Results such as these help formally address between-species brain organization with an aim to improve the translation from preclinical to human results.

---

Much of our knowledge about the human brain is based on knowledge obtained in other species. While numerous species have been used to model the human brain, the mouse has emerged as the most prominent of these, due to its rapid life cycle, straightforward husbandry, and amenability to genetic engineering (Dietrich et al., 2014). The overall assumption in this work is that the knowledge obtained in the mouse ‘model species’ is translatable to the human, due to overall similarities in biological properties of the two species. However, the success rate of such translations have sometimes been disappointing, especially in the case of neuropsychopharmacology (Hay et al., 2014). This is due, in part, to assumptions of between-species similarities not holding (Striedter, 2022). As such, it is becoming increasingly apparent that these assumption should be subjected to explicit empirical validation.

The various divisions of cingulate cortex have repeatedly been shown to be important in many aspects of emotional processing, decision making, and cognitive control (Behrens et al., 2013; Leech and Sharp, 2014; Kolling et al., 2016) and alterations in cingulate morphology are a common observation across a range of psychiatric disorders (Goodkind et al., 2015; Opel et al., 2020). Cingulate cortex is thought to be an evolutionary conserved region in mammals. Indeed, an analysis of common areas across six major mammalian clades suggests that cingulate cortex is present in all and that it could have been part of a limited set of neocortical regions present in early mammals (Kaas, 2011). The combination of common alterations in disease and apparent evolutionary conservation make cingulate cortex an important target for translational neuroscience research.

However, the similarity of rodent and human cingulate has been called into question on a number of grounds. First, it has been argued by some authors that rodent cingulate cortex has organizational features or at least performs functions that are homologous to those of primate dorsolateral prefrontal cortex (Brown and Bowman, 2002; Uylings et al., 2003; Carlén, 2017). Second, among researchers who reject these claims, there still is some debate about how rodent cingulate should be subdivided and how its organization relates to that of the primate (Laubach et al., 2018; van Heukelum et al., 2020). Third, even if cingulate cortex were found to be fully homologous in mouse and human it would be embedded within the larger prefrontal network within the human brain compared to other species (Schaeffer et al., 2020). These arguments continue to be reassessed with the appearance of new data types that enable better and more complete comparisons across species.

One way to explore similarities and differences in brain organization across species is by studying connectivity. The connections of brain areas constitute a unique ‘fingerprint’ and provide information about their incoming information and the influence they exert on other parts of the network (Passingham et al., 2002; Mars et al., 2018). We have previously employed functional connectivity as assessed using resting state fMRI to compare connectivity across humans and non-human primates (Mars et al., 2011, 2016) and humans and mice (Balsters et al., 2020). Cingulate connectivity has been studied using neuroimaging in a number of studies using both diffusion MRI tractography (Beckmann et al., 2009; Smith et al., 2018) and functional connectivity (Hutchison et al., 2012; Schaeffer et al., 2020). Here, we study mouse cingulate functional connectivity, assessing it against the ‘gold standard’ of tracer-based structural connectivity, and compare it with similar data from the human. The goal of the study is to assess to what extent the general organizational principles of cingulate organization are comparable across the two species.

## Materials and methods

### Data-driven analysis of mouse tracer data

We first performed a data-driven parcellation of the rodent cingulate cortex based on structural connectivity as established using tracers following the strategy of Mandino and colleagues (2022). This serves as a baseline for the subsequent analyses using functional MRI data.

We downloaded data from 498 tracer experiments from the independent tracer-based connectivity dataset of the Allen Institute (Oh et al., 2014) using a custom interface. In these experiments C57BL/6 male mice received a viral anterograde tracer injection in various subcortical and cortical sites of the right hemisphere. This viral tracer initiates the coding of a fluorescent protein which accumulates in the axons of neurons. Through visualising this fluorescence, a detailed description of the structural connections between the site of injection and the rest of the brain can be created.

After downloading these tracer experiments, 2000 seeds were placed at even intervals along a region of interest spanning the left hemisphere anterior cingulate area, infralimbic area, prelimbic area, and retrosplenial area (hence referred to as the ‘cingulate ROI’) according to the nomenclature of the Allen mouse brain reference atlas (Wang et al., 2020). Where possible, we will use the terminology of Vogt and Paxinos (2014) for the cingulate and Paxinos and Franklin (2019) for the rest of the brain when discussing our results.

Subsequently, we extracted the tracer-density in these seeds and correlated to the tracer-density recorded in the rest of the brain, thus resulting in a seeds by whole-brain correlation map. The left hemisphere was selected for the seed locations to extract axonal projections, as opposed to cell body-related trace-density. This allowed us to generate connectivity maps based on axonal projection similarity.

Having created the correlation maps, we grouped seed voxels together as a function of their connectivity profiles by performing an independent component analysis (ICA) on the correlations maps using FSL’s *melodic* (Beckmann and Smith, 2005). An independent component provides a spatial map of voxels that have similar correlations to the cingulate seed voxels. Thus, ICA essentially divides the brain, including the cingulate cortex itself, into components based upon their connections with the cingulate cortex. We ran ICA multiple times, each time requesting a different number of components, ranging from four to nine. In general, the components remained stable for the different amount levels of granularity. However more subtle effects become apparent at greater granularities.

### Mouse structural connectivity fingerprints

To summarize the connectivity of the different parts of the cingulate ROI with the rest of the brain, we describe the correlation of connectivity of seed areas in the cingulate with target areas in the rest of the brain as ‘connectivity fingerprints’ (cf. Passingham et al. (2002); Mars et al. (2018)). To this end, we placed ten seeds in the cingulate ROI at even intervals along the rostral-caudal axis. Anteriorly, one seed was placed in area 25 and another one in area 32. More caudally, four seeds were placed in area 24 and 24’ of the mid-cingulate. Finally, we placed an additional four seeds in the retrosplenial area; one of which was placed in the most posterior-lateral part of RSA, just above the post-subiculum.

We chose target regions on the basis of multiple criteria. Firstly, regions were selected that receive projections from, or project to, the cingulate according to existing literature. Secondly, the results from the ICA were used to ensure that the target regions would be able to differentiate between the seeds. For instance, a nucleus of the thalamus that projects strongly to the entire cingulate ROI is of little use for distinguishing between the different cingulate subregions. Finally, target regions were only considered if their homology across mouse and human brains is well established. By these criteria, the following 11 target regions were selected for the mouse connectivity fingerprints: (1) hippocampal formation, (2) amygdala, (3) nucleus accumbens, (4) hypothalamus, (5) caudoputamen, (6) superior colliculus, (7) primary visual area (V1M), (8) secondary motor area, (9) medial parietal association cortex, (10) dorsal agranular insular cortex, (11) ventral orbital cortex.

### Mouse resting state fMRI data

To compare the organization of the cingulate across species, it is preferable to use the same type of data (Mars et al., 2021). We obtained publicly available resting state functional MRI data from both mice and humans. We used these data to estimate ‘functional connectivity’, i.e., similarity in time courses of spontaneous blood oxygenation level dependant contrast fluctuations across voxels.

Mouse resting state fMRI scans were downloaded from an existing pre-processed dataset collection (https://doi.org/10.34973/1he1-5c70) (Grandjean, 2020). For these datasets, mouse functional MRI acquisitions were conducted in accordance with the Swiss federal guidelines and under a license from the Zurich Cantonal Veterinary Office (149/2015) as well as the ethical standards of the Institutional Animal care and use Committee (A*STAR Biological Resource Centre, Singapore). Scans from 188 (106 male, 82 female) healthy wildtype mice (C57BL/6 strain) were downloaded. These scans, in turn, belong to different sub-datasets (Table 1).

**Table 1.**
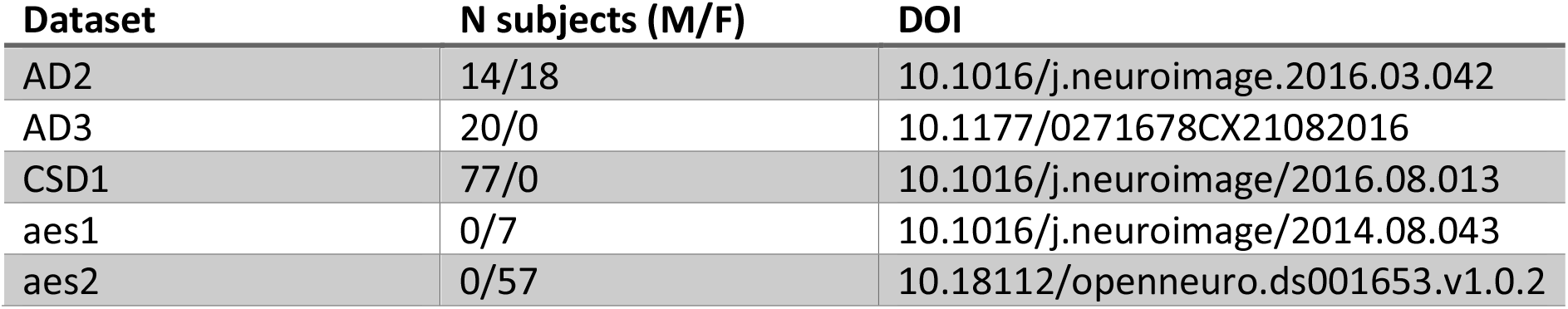
rs-fMRI mouse datasets.

In all these sub-datasets, the mice were anesthetised with isoflurane (see Grandjean et al. (2014) for a detailed protocol). Subsequently, the mice were mechanically ventilated and placed on an MRI-compatible cradle. During the scanning, the anaesthesia was maintained with a combination of isoflurane (0.5%) and medetomidine infusion (0.1 mg/kg/h). Different scanner settings were used for each of the datasets.

The mouse scans were preprocessed as described in Huntenburg et al. (2020). Briefly, the anatomical scans were corrected for the B1-field inhomogeneity, denoised, brain-masked and, via the study template, registered to the Allen reference template (resampled to a 0.2 mm^3^ resolution). The functional scans were despiked, motion corrected, corrected for the B1 field, denoised, brain masked and registered to their anatomical images. Finally, they were bandpass filtered (0.01–0.25 Hz) and an ICA was applied to determine nuisance components which were subsequently filtered out (https://github.com/grandjeanlab/MouseMRIPrep).

The mean timeseries were extracted from the seed and target locations for each scan using *fslmeants* and correlations between the timeseries was calculated for each subject using *1ddot*. Finally, the correlations were averaged over the subjects and visualised into a fingerprint which indicates how a single target regions connects to the different cingulate seeds.

### Human resting state fMRI data

To study human functional connectivity, we used the S1200 7T rs-fMRI dataset of the Human Connectome Project (HCP, available at https://db.humanconnectome.org) (Van Essen et al., 2013). The precise parameters for both the data acquisition and preprocessing have been described elsewhere (Glasser et al., 2013; Smith et al., 2013). In short, rs-fMRI was acquired using a gradient-echo EPI sequence at 7T with the following parameters: TR=1000 ms, TE=22.2 ms, multiband-factor=5, isotropic resolution=1.6 mm, field-of-view= 208*208 mm, bandwidth=1924 Hz/px, Image Acceleration factor=2. Subjects were scanned four times, each session lasted approximately 16 minutes during which 900 volumes were acquired. For the current research, only the first of these scanning sessions with posterior-anterior phase encoding was used.

We carried out quality control based on connectivity plausibility in the S1200 7T dataset (Grandjean et al., 2023). In the end 127 scans were downloaded from the HCP (39 male, 88 female). The scans were already preprocessed with the HCP pipeline (Griffanti et al., 2014; Salimi-Khorshidi et al., 2014) and were further preprocessed by smoothing and bandpass filtering (0.01-0.1 hz) the scans using AFNI 3dTproject. The mean timeseries were extracted for the seed and target locations and correlated to each other using *fslmeants* and AFNI’s *1ddot*, respectively, for each subject. As for the mouse, the correlations were averaged over the subjects and visualised as a single fingerprint for each target region.

To create the human connectivity fingerprints analogous to those in the mouse, seeds were placed in the cingulate along the rostral-caudal axis at even intervals. We used the atlases of Neubert et al. (2015) and Beckmann et al. (2009) to assign approximate area names for anterior and midcingulate and for posterior cingulate and retrosplenial cortex, respectively. Twelve seeds were placed at the following MNI [x y z] coordinates: seed 1 [4 18 −10] (area 25), seed 2 [4 30 −6] (ventral border of area 24 and dorsal border of area 14m), seed 3 [4 40 0] in area 24, seed 4 [4 38 12] (area 24), seed 5 [4 30 22] (area 24), seed 6 [4 16 32] (border of area 24 and RCZa), seed 7 [4 0 38] (area 23ab/RCZp), seed 8 [4 −14 36] (area 23ab/RCZp), seed 9 [4 −30 38] (area 23ab/RCZp), seed 10 [4 −40 32] (area 31/23ab), seed 11 [4 −48 16] (area 23ab), seed 12 [10 −46 8] (area 23ab).

Subsequently, twelve target regions were selected which are considered to be homologues to the mouse target regions, namely: the hippocampal formation [26 −16 −20] (according to Amunts et al. (2000), the amygdala [22 −6 −16] (Amunts et al., 2005), the nucleus accumbens [10 16 −4], the hypothalamus [4 −6 −8], the caudate nucleus [12 16 4], the putamen [12 6 4], the superior colliculus [6 −32 −4], the primary visual area [10 −94 −2] (V1; Amunts et al. (2000)), the supplementary motor area [8 −4 60] (Neubert et al., 2015), the superior parietal lobule [30 −56 62] (SPLC; Mars et al. (2011)), anterior insula [42 12 −6], and orbitofrontal cortex [6 30 −20] (area 14m; Neubert et al. (2015)).

For a follow-up analysis to investigate connectivity of the cingulate seeds with human granular prefrontal areas, we placed additional seeds in granular orbitofrontal cortex [4 46 - 12] (area 11m; Neubert et al. (2015), medial frontal pole [6 62 4] (FPm; Neubert et al. (2015)), lateral frontal pole [25 57 5] (FPl; Neubert et al. (2014)), medial frontal gyrus [28 40 32] (area 9/46D; Sallet et al. (2013), and area 9m [8 58 28] (Neubert et al., 2015).

To determine whether cingulate seeds possessed characteristic patterns of connectivity probability from the target areas considered, we carried out repeated-measures analyses of variance on the data, with factors for seed and target area. Huyhn-Feldt adjustment was applied where necessary.

### Data and materials availability

The Allen Institute tracer data is available for non-commercial purpose (http://connectivity.brain-map.org/). The mouse resting-state fMRI data is available under the terms of the CC-BY-4.0 licence (https://doi.org/10.34973/1he1-5c70). The human resting-state fMRI data is available under the terms of the HCP Data Use terms (https://humanconnectome.org/).

## Results

### Mouse tracer data

We set out to examine the structural networks of the cingulate area in the mouse. Earlier studies demonstrated that projection similarity across a viral tracer dataset can be used to examine the projectome of seed regions (Mandino et al., 2021). Here, we applied the same method by sampling 2000 seeds across the cingulate cortex. To summarize the outcomes of the seed-based maps, we applied an independent component analysis. In general, the components of the ICA remained stable for the different levels of granularity, although more subtle effects become apparent at greater granularities. We here present the solution for six components, to provide a balance between granularity and coarseness. Cluster solutions for n=4 and 9 components are presented in the supplementary material.

The first component overlapped with the anterior part of our ROI, in the territory delineated as area 25 and partly with area 32 by Vogt and Paxinos (2014) (Figure 1, panel A). Outside the cingulate this component overlapped with limbic structures such as the hippocampal formation, amygdala, and the nucleus accumbens. Another, weaker, orbitofrontal component (Figure 1, panel F) was component mostly associated with the anterior cingulate areas, but did not show much association outside the ROI.

**Figure 1.**
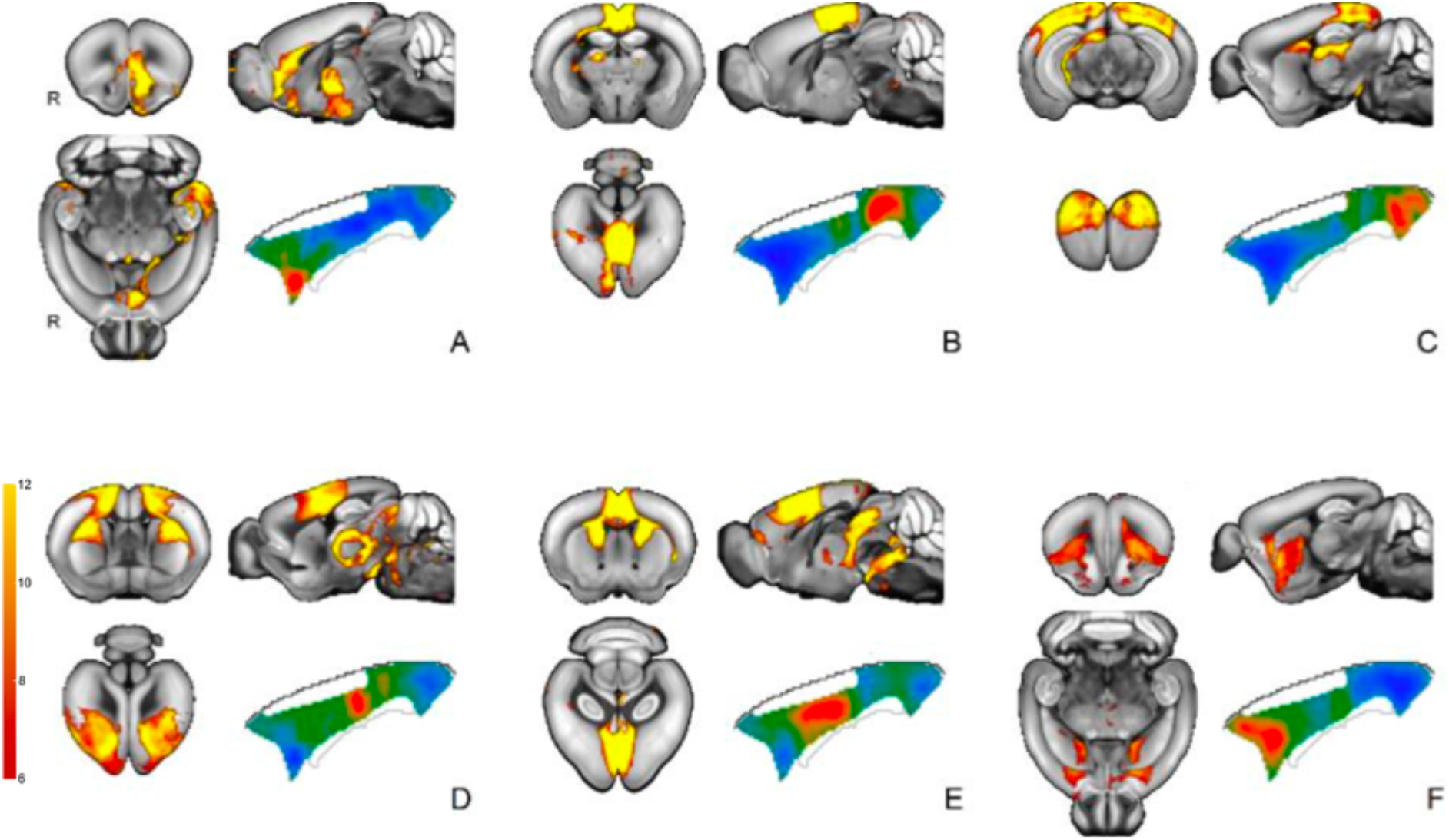
Independent component analysis of mouse structural connectivity from viral anterograde tracer injection studies relative to the cingulate area. Six component solution depicting the spatial maps (hot colors) and the cingulate area seeds associated with the component (blue to red) projected onto the outline of the cingulate area.

At the posterior end of the ROI, a component overlapped with the retrosplenial area (Figure 1, panel B). This component was relatively self-contained and did not extend to as many structures apart from the superior colliculus and the midbrain. In addition, there is a component that overlaps both with parts of the retrosplenial area and which also overlaps with visual regions such as the primary visual areas, the lateral geniculate complex, and the superior colliculus (Figure 1, panel C).

Two components showed strong overlap with the mid part of the cingulate, overlapping with the territory of areas 24 and 24’ (Figure 1, panel D/E). The first of these components also overlapped with the periaqueductal grey, the pons, the cerebral peduncles, and various nuclei of the thalamus. The other component also overlapped with parts of the caudoputamen, periaqueductal grey, the thalamus and pons (nucleus raphe).

In sum, the data-driven decomposition from the tracer studies identifies components mostly organized along the anterior-posterior axis. Anterior components showed overlap with amygdala and nucleus accumbens, among other areas. Mid-cingulate areas overlapped with caudate and secondary motor cortex, posterior areas overlapped with visual and hippocampal structures. To confirm these results and allow more direct comparisons across the length of the ROI, we placed ten seeds spread across the anterior to posterior dimension. For each seed, we established the whole-brain tracer connectivity and correlated that with the connectivity of a series of target areas. This allows a more direct comparison across different parts of the cingulate as well as a comparison with the resting state fMRI data described below.

The connectivity fingerprints recapitulated the observations from the ICA. Specifically, anterior seeds tended to show high connectivity with amygdala and nucleus accumbens targets (Figure 2). Orbitofrontal cortex connectivity was also mostly associated with anterior cingulate seeds, but more widespread. Hippocampus and hypothalamus both reach the most anterior seed, with the hippocampus also showing strong connectivity with the most posterior seed in the retrosplenial area.

**Figure 2.**
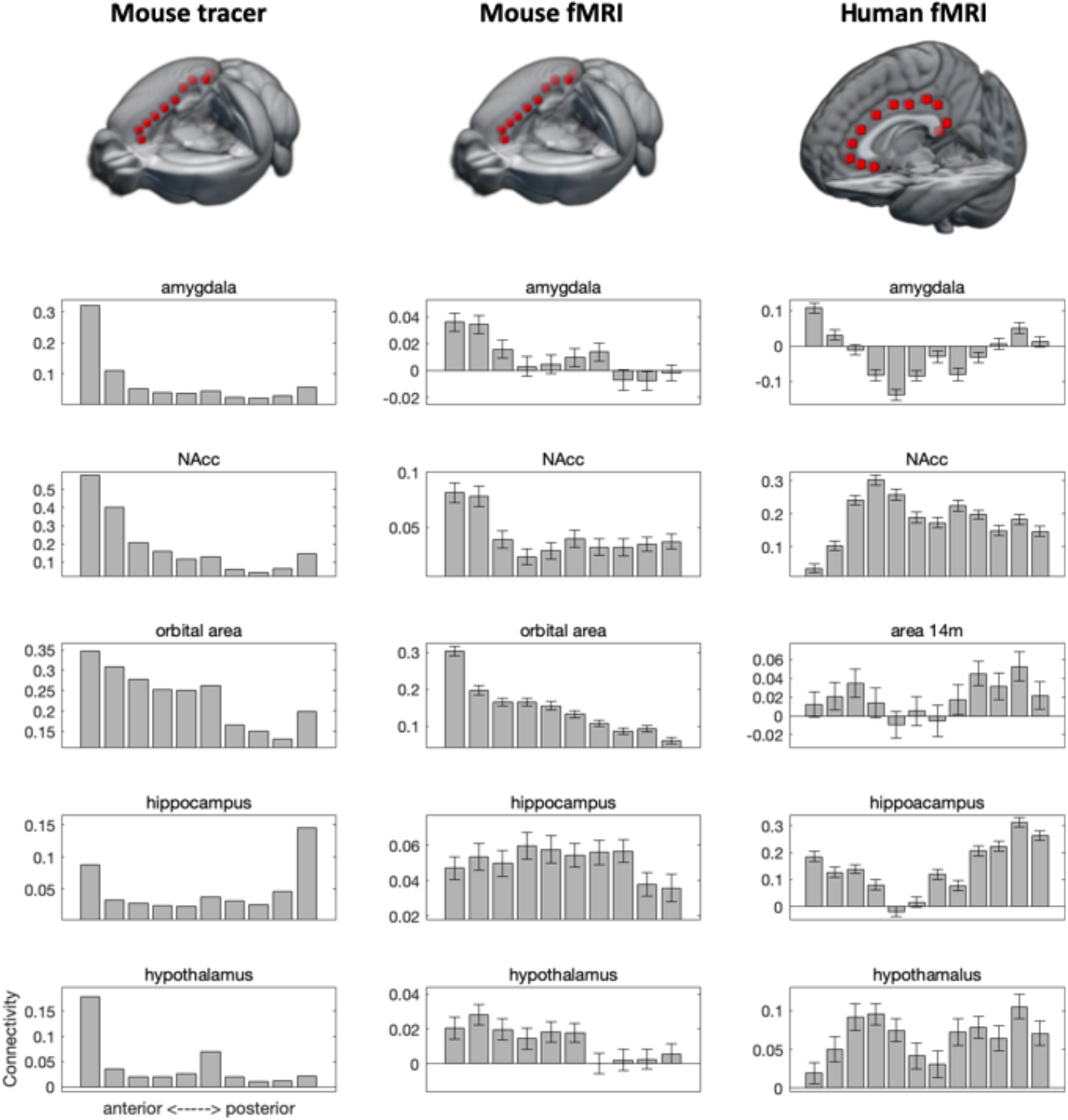
Connectivity of subcortical and orbitofrontal target areas with cingulate seed areas in all modalities and species. Error bars indicate +/- SEM.

**Figure 3.**
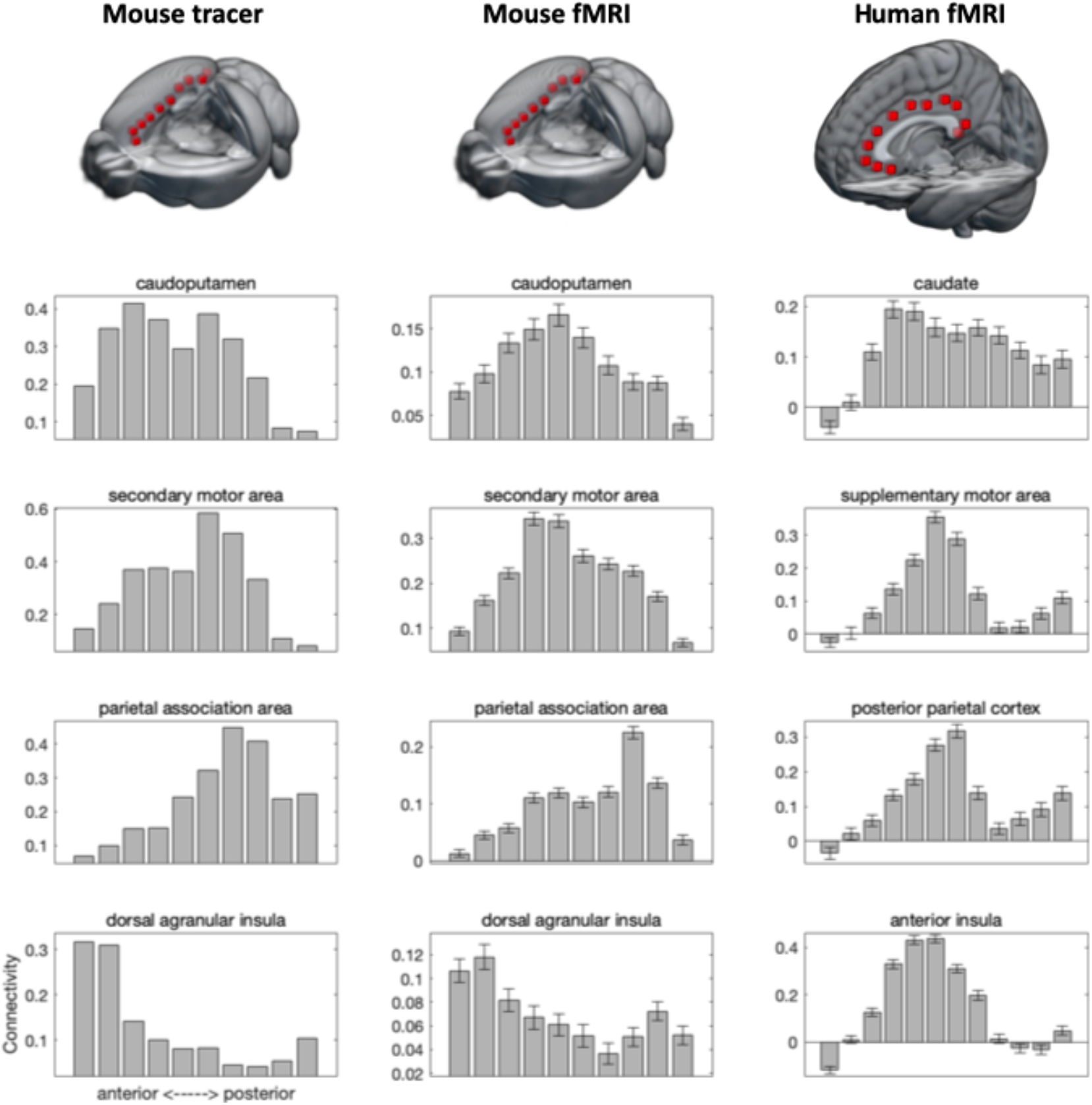
Connectivity of caudate, motor and parietal, and insula target areas with cingulate seed areas in all modalities and species. Error bars indicate +/- SEM.

Again in accordance with the ICA decomposition results, caudoputamen and secondary motor cortex showed strong connectivity with medial seeds, with caudoputamen connectivity a bit more widespread than that of secondary motor cortex. The medial parietal association area followed a pattern similar to that of secondary motor cortex. Finally, we looked at the connectivity of the anterior insula. This showed a quite confined connectivity with the two most anterior seeds.

### Mouse and human resting state fMRI data

To enable comparison of cingulate connectivity across the mouse and the human, we analysed resting state functional MRI data. This allows us to assess the similarity in the time courses of spontaneous activation of all areas of the brain. Such ‘functional connectivity’ is not the same as the anatomical connectivity assessed using tracers, but the two have been shown to correlate (Grandjean et al., 2017). Using functional connectivity allows us to compare human and mouse brain organization using the same method. We examined functional connectivity, using MRI data, of the mouse seed areas in cingulate cortex with the same targets examined in the tracer data set. We then compared the MRI-based estimates of cingulate connectivity in the mouse to connectivity of twelve seeds placed in anterior-to-posterior locations and the homologous target areas. For both the mouse (*F* = 59.409, *p* < 0.001) and the human (*F* = 53.050, *p* < 0.001), functionally connectivity showed a seed by target interaction, indicating that the target areas can be used to distinguish between different cingulate seeds.

Functional connectivity of mouse amygdala, nucleus accumbens, and orbitofrontal cortex followed a pattern very similar to that of the tracer data, with connectivity mostly restricted to the anterior seeds. The same was true for human amygdala and to a lesser extent orbitofrontal cortex, which also showed some more posterior connectivity. Nucleus accumbens, in contrast, has a more widespread connectivity pattern in the human. As was the case in the mouse tracer data set, human hippocampus showed functional connectivity with the anterior and posterior, but not mid, cingulate seeds, although the human pattern is more widespread than the mouse tracer. Mouse hippocampal functional connectivity, by contrast, showed a very widespread pattern of functional connectivity. Hypothalamic connectivity was also more widespread in functional data than in tracer data.

We next investigated a number of target areas that in the human are known to show connectivity mostly with the mid part of the cingulate, including the territory of the cingulate motor areas. Mouse caudoputamen and human caudate both showed a widespread connectivity with the cingulate seeds, but mostly peaking in midcingulate areas. The pattern was much more clearcut in the cases of connectivity with the human supplementary motor area and posterior parietal cortex; these areas had a clear peak of connectivity with midcingulate areas. We also observed a higher functional connectivity of the mouse secondary motor area and parietal association area with midcingulate areas, although the patterns was much less clear than in the human and the peaks of the two target areas did not overlap. This difference in pattern was apparent in both the mouse tracer and resting state functional MRI data. The biggest difference between mouse and human was observed in connectivity with the insular target area. In the human, insula showed a clear affinity with the midcingulate seeds, but in the mouse both the tracer and functional data showed strongest connectivity with anterior, and to a lesser extent posterior, cingulate seed areas.

### Human prefrontal connectivity

As a follow-up, we investigated the connectivity of human prefrontal areas with the cingulate seeds. Granular tissue of the sort found in human prefrontal cortex is not found in rodents (Preuss and Wise, 2022). It is therefore important to quantify how our region of interest, the cingulate cortex, is connected to it in order to understand any claims of similarity or difference across species. In addition to the agranular area 14m described above, we quantified functional connectivity of the cingulate seeds with granular orbital area 11m, medial area 9, dorsolateral area 9/46D, and the medial (FPm) and lateral frontal pole (FPl).

All of these regions showed at least some functional connectivity to at least some of the cingulate regions, although there were marked differences in the profile of connections (*F* = 25.130, *p* < 0.001) (Figure 4). Medial prefrontal areas tended to show stronger connectivity with the most anterior and posterior cingulate seeds. In contrast, lateral 9/46D shows strongest connectivity with the mid part of the cingulate. The lateral frontal pole provided a mixture, with strongest connectivity anterior and posterior cingulate cortex, but noticeably also with mid-level cingulate cortex. In sum, all frontal areas tested showed a positive functional connectivity with human cingulate cortex.

**Figure 4.**
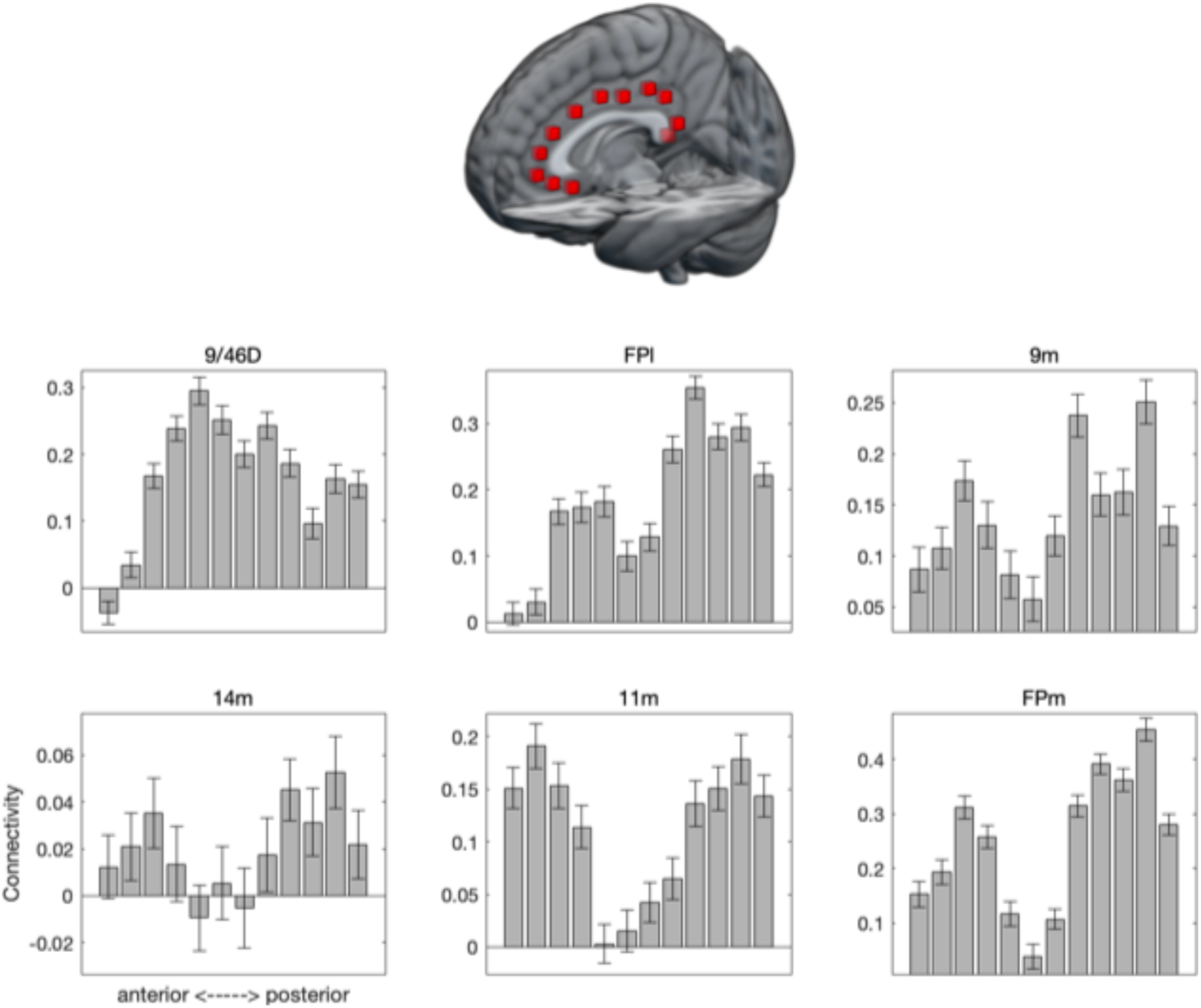
Human functional connectivity of all cingulate seed areas with six prefrontal areas. Error bars indicate +/- SEM.

## Discussion

We set out to investigate whether the mouse and human cingulate cortex are organized according to similar principles in terms of their connectivity to other parts of the brain. Overall, we show that the two species’ cingulate cortices follow broadly similar principles, with anterior areas mostly interacting with amygdala, nucleus accumbens, and orbitofrontal cortex; a midcingulate territory interacting with premotor and posterior parietal cortex; and a retrosplenial zone interacting with hippocampus. The similarity of these patterns is inconsistent with theories ascribing homology of rodent anterior cingulate with primate granular prefrontal cortex (Krettek and Price, 1977; Eichenbaum et al., 1983) or that suggest rodent cingulate contains a mixture of primate cingulate and granular prefrontal features (Uylings et al., 2003). Rather, it is consistent with notions that infralimbic cortex in the mouse is similar to primate area 25 (Alexander et al., 2019), that there is a midcingulate zone with parietal and premotor connections in both species (Vogt, 2016), and a generally similar anterior-posterior organization in both species (cf. Van Heukelum et al. (2020)).

These similarities between species notwithstanding, some differences are apparent. Overall, many human cingulate areas have connectivity profiles that seem more distinct from one another that do those of the mouse cingulate cortex. Earlier work suggested parietal connectivity in the mouse is with both midcingulate and retrosplenial cortex (Zingg et al., 2014) and we indeed see rather widespread parietal connectivity in the mouse. Premotor and parietal connectivity are more restricted to the mid-cingulate cortex in the human. Human midcingulate cortex is thought to contain distinct anterior and posterior subdivisions (Vogt, 2016), the first of which is not present in the mouse (Vogt and Paxinos, 2014). In the human, parietal connectivity is stronger in the posterior part of midcingulate.

Mouse connectivity as assessed using tracers and using resting state functional MRI were overall in agreement, but some differences were noticeable. Hypothalamic connectivity as assessed using tracers was very strong in the most anterior parts of the cingulate, consistent with earlier reports (van Heukelum et al., 2020), but functional connectivity was more broadly distributed in both species. Human hippocampal functional connectivity resembled that of mouse tracers, but not mouse functional connectivity. This could be due to the effect of anaesthesia on the resting state fMRI of the mouse, but this awaits systematic comparison. Hippocampal and hypothalamic connectivity to posterior seeds was much stronger in the human than in the mouse. This is potentially due to the presence of a large posterior cingulate in the human (Bzdok et al., 2015), whereas in the mouse this area only contains a retrosplenial cortex (Vogt and Paxinos, 2014).

The most clear dissociation between the mouse and human data was in the connectivity of the insula. This is true even though we seeded in territory commonly described as agranular anterior insula in both species. In the mouse, the insula seed showed connectivity with anterior parts of the cingulate in both tracer and rs-fMRI data, while in the human the insula seed showed strong functional connectivity with midcingulate areas. The human results are in accordance with models of dorsal anterior cingulate function in cognitive control and the participation of the two regions in a so-called salience network (Seeley et al., 2007; Menon, 2011). Previous work has shown that the salience network, although present in both species, has different associations with the serotonergic network across human and mouse (Mandino et al., 2022), suggesting that the area has changed substantially since the last common ancestor of mice and humans. Alternatively, the insula seed areas we selected in human and mouse are not homologous. We have taken a region commonly used in neuroimaging studies as our human anterior insula (Cieslik et al., 2015; Molnar-Szakacs and Uddin, 2022), but Ongur and Price (2000) describe a number of insular regions more anteriorly, on the caudal orbital surface. Whether the anatomical similarity between these human areas and mouse insular areas is greater than that between our human insula seed and the mouse is a topic for further investigation.

In general, it is important that the target areas used are homologous across species when comparing connectivity across species. Here, we have taken care to use regions that are identified as such, but some discussion is in order especially when in case of targets in the neocortex. The approach used here can be used to ascertain the degree of similarity/difference between any areas in human and mouse. With respect to premotor cortex, the human brain contains areas that have no homolog in the rodent (Wise, 2006), but the two brains’ premotor cortices do follow largely similar organizational principles (Lazari et al., 2023). The human ventral frontal cortex contains agranular, dysgranular, and granular areas, but rodent prefrontal cortex contains only agranular areas (Wise, 2008; Rudebeck and Izquierdo, 2022). We here used human area 14m as defined by Neubert et al. (2015), which is posterior to the granular areas, as our orbitofrontal cortex seeds. We do note that similar results could we obtained using targets in granular area 11m. Posterior parietal cortex is dramatically expanded in primates compared to other mammals (Krubitzer and Padberg, 2009), but a mouse parietal association area that is homologous to primate posterior parietal cortex has been identified (Lyamzin and Benucci, 2019). The current human parietal results are similar for targets overlapping with human MIP or 7A (Mars et al., 2011).

Apart from these differences described earlier, it should be taken into account that the human cingulate is embedded within a much larger and more elaborate neocortical network than that of the mouse. This means that, even if the overall organization of the two species’ cingulate with homologous areas is comparable, connectivity with non-homologous areas mean that the overall connectivity profile can still be quite distinct. This was previously shown to be the case for the human dorsal caudate; although striatal connectivity follows similar organisational principles in both species, connectivity of the dorsal caudate with the human frontal pole means that its connectivity profile is distinct from any found in the mouse (Balsters et al., 2020). Connectivity between the human medial frontal gyrus and human cingulate is evident in the present data as in previous studies (Sallet et al., 2013; Loh et al., 2018). In the striatum, areas with a distinct human connectivity profile were associated with higher-order cognitive processes, including executive control and language. It remains to be seen whether functional differences are found between the two species’ cingulate regions.

Model species are an essential part of research in biology and by extension neuroscience (Striedter, 2022). Differences between the model and the species of ultimate interest, i.e., the human, are to be expected and do not necessarily present a problem for translational neuroscience, as long as these differences are properly understood. Whole-brain, high-throughput data are now increasingly available and allow us to gain a much more systematic understanding of such differences than ever before (Mars et al., 2014). The present work contributes to this effort by comparing a major target area for clinical research across species by means of connectivity. Future work will focus on comparing these results obtained using comparative connectivity with those obtained using other modalities, such as spatial patterns of gene expression, tissue properties, and receptor densities (Vogt et al., 2013; Beauchamp et al., 2022).

In sum, this work shows the feasibility of extending existing approaches of comparing frontal cortical organization across species using functional MRI to rodent-human comparisons. The results show a generally conserved macro-level organization, although there are important differences in both regional specialization and embedding within larger cortical networks. Such differences are important to take into account when performing between-species translations in the context of clinically relevant research.

